# Disruption of CaMKII biomolecular condensation induces attention-deficit/hyperactivity disorder-like phenotypes

**DOI:** 10.64898/2026.06.24.734155

**Authors:** Yui Sugiyama, Chisato Suematsu, Risa Yamada, Atsuki Goda, Shogo Miyata, Tokiwa Yamasaki, Ryuichi Shigemoto, Anjuli Dijkmans, Danielle C. M. Veenma, Shoji Takada, Geeske M. van Woerden, Yasunori Hayashi, Takeo Saneyoshi

## Abstract

CaMKII is a multifunctional kinase essential for synaptic plasticity and memory formation. While its canonical role involves enzymatic phosphorylation, recent evidence suggests CaMKII also functions through liquid–liquid phase separation (LLPS) with substrate proteins, including GluN2B-containing NMDA receptors. However, the physiological significance remains unclear. Here, we generated CaMKII α subtype knock-in (KI) mice carrying a point mutation (I205K) in the hydrophobic pocket, a key interface required for LLPS. These mice exhibited a complete loss of structural long-term potentiation (sLTP) despite normal spine morphology, marked hyperactivity and profound deficits in aversive memory formation. Atomoxetine, an approved attention-deficit/hyperactivity disorder (ADHD) treatment, ameliorated the hyperactive phenotype. Notably, we identified a patient carrying the I205N variant presenting with ADHD and mild intellectual disability, mirroring the behavioral features observed in I205K KI mice. Additional neurodevelopmental disorder-associated variants within the same hydrophobic pocket similarly disrupted LLPS *in vitro*. Molecular dynamics simulations revealed these variants destabilize the CaMKIIα–GluN2B interaction through distinct mechanisms that perturb the dynamic stability of the binding interface. These findings establish CaMKIIα-mediated phase separation as critical for linking synaptic molecular assembly to cognitive function and provide a unifying molecular basis for synaptic disorganization, hyperactivity, and memory deficits associated with neurodevelopmental disorders.

**Significance:** CaMKIIα is essential for synaptic plasticity and memory, yet the contribution of its non-catalytic functions, including liquid–liquid phase separation (LLPS), to brain function remains unclear. Here, we show that disrupting CaMKIIα LLPS with its synaptic partners impairs synaptic localization, abolishes LTP, and causes profound memory deficits in vivo. A knock-in mouse carrying the I205K mutation showed hyperactivity and impaired learning. This recapitulate the clinical phenotypes of a human patients with the I205N variant, which is associated with ADHD and intellectual disability. Together, these findings establish LLPS as a critical mechanism underlying CaMKIIα function in the brain and provide a disease-relevant framework for understanding synaptic dysfunction in neurodevelopmental disorders.

## Introduction

The Ca^2+^/calmodulin-dependent protein kinase II (CaMKII) is a pivotal signaling molecule in the central nervous system, essential for activity-dependent synaptic plasticity, learning, and memory ^1,2^. CaMKII has a unique biochemical property—autophosphorylation-driven, Ca^2+^/calmodulin-independent kinase activity—that enables sustained substrate phosphorylation following transient calcium influx ^2^. Beyond its enzymatic role, CaMKII also functions as a structural protein through dynamic interactions that localize it to postsynaptic compartments, particularly in response to NMDA receptor activation ^3–6^.

Traditionally, CaMKII activation during long-term potentiation (LTP) has been understood as a linear cascade: calcium influx through NMDA receptors activates CaMKII, leading to autophosphorylation at T286 and generating Ca^2+^-independent “autonomous” kinase activity, which has been proposed to encode molecular memory traces ^7,8^. However, this classical view has been challenged. Recent evidence suggests that kinase activity is not required for the maintenance of LTP, but the structural roles mediated by the interaction of CaMKII with the GluN2B subunit of the NMDA receptor play a critical role in synaptic plasticity ^9–11^. Nevertheless, knock-in mice harboring GluN2B mutations that disrupt CaMKII binding exhibit only mild learning impairments ^12^, implying the existence of additional molecular mechanisms underlying memory encoding. Indeed, we and others have identified that a CaMKII inhibitor peptide, Rac-guanine nucleotide exchange factor Tiam1, densin-180/LRRC7, eag K^+^-channel, and Ras-like GTPase Rem2 can also interact with the same site of CaMKII ^5,13,14^.

Through interactions with these proteins, CaMKII can undergo liquid–liquid phase separation (LLPS), forming dynamic condensates that organize the postsynaptic signaling complex ^15,16^. This emerging concept suggests that CaMKII LLPS may act as a biophysical mechanism for molecular clustering and signal integration at synapses. Yet, the physiological relevance of CaMKII LLPS in the intact brain and behavior remains largely unexplored.

To address this, we generated a CaMKII α subunit knock-in mice carrying a mutation that selectively abolishes LLPS. I205 lies within the hydrophobic pocket that mediates interactions with proteins such as GluN2B and Tiam1^13^. We found that the I205K mutation abolished LLPS *in vitro* without measurably affecting kinase activity ^16^. In the brain, this mutation disrupted CaMKII-GluN2B co-localization and synaptic enrichment, while the basal expression and phosphorylation levels of CaMKII as well as glutamate receptor distribution were only modestly affected. In contrast, structural LTP (sLTP) induced by glutamate uncaging was significantly impaired, indicating a selective requirement of CaMKII LLPS in activity-dependent synapse remodelling. The KI mice exhibited pronounced hyperactivity that was ameliorated by atomoxetine, a therapeutic drug for ADHD, together with severe deficits in aversive learning that could not be rescued by repeated trials. Furthermore, analysis of a human cohort with neurodevelopmental disorders identified two *de novo* CaMKII variants mapping to the same hydrophobic pocket. One patient carrying the I205N variant was diagnosed with ADHD accompanied with mild intellectual disability, supporting the validity of our mouse model. Together, these findings demonstrate that CaMKIIα LLPS is essential for the organization of postsynaptic signaling and for learning and memory. Our study reveals a previously unrecognized mechanism by which disruption of CaMKII LLPS behavior contributes to the pathophysiology of neurodevelopmental disorders such as ADHD and intellectual disability.

## Materials and Methods

### Animals

All procedures were conducted in accordance with the institutional guidelines and protocols approved by Kyoto University Animal Experiment Committee. The CaMKII I205K knock-in mouse line was generated by CRISPR/Cas9-mediated genome editing as previously described ^17^. For genotyping, genomic DNA was extracted from tail tissues using proteinase K digestion, followed by PCR amplification with the following primers:

Forward, 5’-GTTCGCAGGGACACCTGGATAC-3’

Reverse (WT), 5’-GGGGGATACCCAACCAGCAAGA-3’

Reverse (I205K), 5’-GGGGGATACCCAACGAGAAGCT-3’

PCR products were analyzed by agarose gel electrophoresis to distinguish WT and mutant alleles. Male homozygous KI and WT littermates (8–16 weeks old) were used for experiments. In a repeated exposure condition of the inhibitory avoidance test, female mice were also used. The I205K mice displayed aggression toward their cage mates. Moreover, although reproductive behaviors appeared normal, abnormalities in nursing behavior were observed. Because of these issues, it was difficult to obtain a sufficient number of I205K mice for behavioral testing through natural breeding. Therefore, the offspring were derived via in vitro fertilization and nursed by foster mothers. Mice were maintained on a 12-hr light/dark cycle (lights on at 8:00 am) with *ad libitum* access to food and water.

### Antibodies

Primary Antibodies: Anti-CaMKIIα mouse monoclonal antibody (clone 6G9, Sc-32288) was obtained from Santa Cruz Biotechnology; anti-NR2B rabbit polyclonal antibody (06-600), and anti-GluN1 mouse monoclonal antibody (MAB363) from Millipore; anti-Tiam1 sheep antibody from R&D; anti-FLAG M2 mouse monoclonal antibody from Sigma; anti-mouse IgG CF633 (20124) from Biotium; anti-rabbit IgG Cy3 (711-165-152) from Jackson ImmunoResearch Laboratories Inc; phospho-specific antibody against PKA substrate (#9621) from Cell Signaling Technologies. Anti-GluA1-3 rabbit polyclonal antibody was described previously ^18^.

### Immunostaining and colocalization analysis of CaMKII and GluN2B

Mice were intracardially fixed with 4% paraformaldehyde and sectioned at 50 µm thickness with a vibratome. The sections were treated with pepsin (1 mg/ml 15 min at 37°C) for antigen retrieval and then incubated with primary antibodies diluted (1:250) in blocking buffer (Tris-buffered saline (TBS, 25 mM Tris-HCl, 137 mM NaCl, 2.7 mM KCl, pH 7.5) supplemented with 0.5 % fish gelatine, 0.1 % TritonX-100, 0.02 % NaN_3_) at room temperature for 16 hours. The slices were then stained with an appropriate secondary antibody and counterstained with Hoechst 33342. Images were acquired using a confocal microscope (LSM780 + Airyscan, ZEISS Microscopy). For quantification of colocalization, background-subtracted images were analyzed using Pearson’s correlation coefficient and M1 & M2 coefficients, which were calculated with the JA-COP plugin for ImageJ/Fiji ^19^.

### SDS-digested freeze-fracture replica labelling (SDS-FRL)

For each genotype, five mice were used for SDS-digested freeze-fracture replica labelling (SDS-FRL), and two replicas were prepared from each mouse as described previously ^20^. Briefly, mice were anesthetized with isoflurane and perfused with 2% paraformaldehyde (PFA) and 15% saturated picric acid. Following perfusion, the brains were extracted and post-fixed in 2% PFA. Coronal sections (80 µm thickness) were cut, and hippocampal slices containing all layers of the CA1 region were trimmed and cryoprotected in 30% glycerol in phosphate buffer (PB, 0.1 M NaH_2_PO_4_, pH 7.4). Slices were frozen using a high-pressure freezing machine (HPM010; Bal-Tec, Balzers, Liechtenstein), fractured at −117°C, and coated sequentially with 5 nm carbon, 2 nm platinum-carbon and 20 nm carbon using a freeze–fracture machine (BAF060; Bal-Tec). Tissue was then dissolved in SDS-solution (2.5% SDS, 20% sucrose, 15 mM Tris, pH 8.3) with gentle shaking for 18 h at 80°C. Replicas were washed once in SDS solution and three times in washing buffer (0.1% Tween-20, 0.05% BSA, 0.05% NaN_3_ in Tris-buffered saline [TBS]), then incubated in blocking buffer (5% BSA in washing buffer) for 30 min. Replicas were incubated overnight at 15°C with a mixture of rabbit anti-GluA1-3 and mouse anti-GluN1 primary antibodies diluted in blocking buffer. After washing three times with washing buffer, the replicas were incubated again in blocking buffer for 30 min and then in a mixture of secondary antibodies (goat anti-rabbit IgG conjugated to 5 nm gold and goat anti-mouse IgG conjugated to 10 nm gold; for GluA1-3 and GluN1, respectively) diluted in blocking buffer overnight at 15°C. Finally, replicas were washed three times in washing buffer, once in TBS and twice in Milli-Q water and mounted on copper grids for electron microscopy observation.

### Electron Microscopic Imaging and Analysis

Labeled replicas were imaged using a Tecnai 10 (FEI, 80kV accelerating voltage) or Tecnai 12 electron microscope (FEI, 120kV accelerating voltage). Images were acquired from the intermediate CA1 region of the hippocampus. During imaging, the sample was tilted to maximize the visible area of the synaptic membrane. Postsynaptic densities (PSDs) were identified based on clusters of intramembrane particles (IMPs) on the exoplasmic face (E-face). Only PSDs with more than 90% of their area contained within the E-face were included in the analysis. Quantitative analyses were performed using *Darea* software ^20,21^. PSDs were manually demarcated as areas containing tightly packed IMP clusters. For the Center–Periphery Index (CPI) analysis, some PSDs were re-demarcated to obtain a smoother contour, representing the presumed complete synaptic profile for more reliable CPI measurement. Gold particles within the demarcated area were detected automatically using the *Darea* software and manually corrected when necessary. The count included particles whose centers were located inside or within 30 nm of the demarcation border.

For each gold particle, the CPI was determined using *Darea* software, and the mean value was calculated for each synapse. CPI is calculated using the following formula:

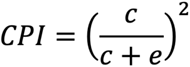

where *c* represents the distance from the gold particle to the center of gravity of the synapse, and *e* represents the distance from the particle to the nearest edge of the demarcated synapse. For particles randomly distributed within a circular area, the expected CPI value is 0.5. For synapses, which are usually not perfectly circular, CPIs between 0.5 and 0.6 are typically observed for randomly distributed particles. Random distribution simulations were performed using a Monte Carlo approach as described previously ^20^. Each simulation was repeated 50 times and averaged for each synapse. Statistical analyses were performed using Prism10 (GraphPad, San Diego, CA, USA) and R (R Project for Statistical. Computing, Vienna, Austria).

### Protein purification and in vitro LLPS assay

Recombinant human CaMKIIα (WT, I205K, I205N, F98S, and E183V) were expressed in Sf9 cells using the DefBac baculovirus system ^22^. The GluN2B cytoplasmic tail (amino acids 1226-1482) and calmodulin were expressed in bacterial strain BL21(DE3) pLysS-codon plus RIL cells (Agilent). All recombinant proteins were purified by affinity chromatography followed by ion-exchange and size exclusion chromatography (Cytiva). The purity of the protein was confirmed by SDS-PAGE. Liquid-liquid phase separation (LLPS) assay was performed as described previously ^16^. Purified proteins (10 µM each) were mixed in a buffer containing 50 mM Tris-HCl (pH 8.0), 100 mM NaCl, 1 mM TCEP, 5 mM MgCl_2_, 2.5 mM ATP, 2 mM CaCl_2_ in the presence of Ca^2+^/calmodulin. Liquid condensate was visualized by confocal microscopy.

### Kinase assays

CaMKII enzymatic activity was measured using autocamtide-2 (AC2, KKALRRQETVDAL) as substrate, which was prepared as GST-fusion protein expressed in bacterial cell as previously described ^5^. Kinase reactions were carried out in a solution containing 50 mM HEPES (pH 7.5), 10 mM MgCl_2_, 2.5 mM DTT, 0.005% Tween20, 50 µM ATP, either in the presence of 5 mM EGTA, or 1 mM CaCl_2_, and 1.5 µM calmodulin. Autophosphorylated CaMKII proteins were generated prior to the kinase reaction in 50 mM HEPES (pH 7.5), 1 mM CaCl_2_, 10 mM MgCl_2_, 2.5 mM DTT, 0.005% Tween20, 1 µM CaMKII, 13.3 µg/µL BSA for 30 min at 30°C, followed by addition of EGTA at 10 mM final concentration. Kinase reactions were 30℃ for 30 minutes. Reaction was terminated by adding SDS-PAGE sample buffer and boiled for 5 minutes before subjected to western blotting. Phosphorylation of AC2 was analyzed by western blotting using phospho-specific antibody against PKA substrate.

### Immunoblotting and co-immunoprecipitation

Hippocampal lysates were prepared in lysis buffer containing 50 mM Tris-HCl (pH 8.5), 150 mM NaCl, 1% deoxycholate, 10% glycerol, 1 mM Na_3_VO_4_, 10 mM NaF, 1 mM β-glycerophosphate, 1 µM microcystin LR, 1 × phosphatase inhibitor cocktail (Nacalai Tesque), and 1 × cOmplete Protease Inhibitor Cocktail EDTA free (Roche). Lysates were centrifugated at 100,000 × *g* for 30 min at 4℃, and the resulting supernatant was used for immunoprecipitation. For each reaction, 4 µg of anti-Tiam1 antibody or control sheep IgG was incubated with IgG-Accept Beads (Nacalai Tesque) for 2-4 hours at 4℃ with rotation. The beads were then washed three times with 1 mL of lysis buffer, and bound proteins were eluted with SDS-PAGE sample buffer. Elutes were analyzed by western blotting.

### Golgi staining and electron microscopy

Golgi-Cox staining was performed using the SuperGolgi kit (Bioenno Tech) according to the manufactures’ instructions, as previously described ^17^. Coronal brain sections containing hippocampus were prepared, and pyramidal neurons in the CA1 regions were selected for analysis. Spine density and morphology were quantified from apical dendrites of fully impregnated neurons under a light microscope using FIJI (NIH) ^19^. Only clearly stained and isolated neurons were included in the analysis to avoid overlapping dendritic structure.

### Behavioral tests

Homozygous CaMKIIα I205K knock-in mice and their wild-type littermates were used for all behavioral analyses. Most behavioral analyses were conducted using male mice; however, both male and female mice were used for the multiple-exposure of stimulation condition in the inhibitory avoidance test.

### Grip strength test and wire hang test

Forelimb grip strength was measured using a grip strength meter (O’Hara & Co., Tokyo, Japan). Mice were lifted by the tail and allowed to grasp a wire grid with their forepaws. They were then gently pulled backward until they released the grid. The peak force (Newtons, N) was recorded, and the highest value from three trials was used for analysis. For the wire hang test, each mouse was placed on a wire mesh that was subsequently inverted and gently shaken to encourage gripping. The latency to fall was recorded, with a 60-s cutoff.

### Hot plate test

Thermal nociception was assessed using a hot plate apparatus (Columbus Instruments). Nine- to 12-wk-old mice were placed on a plate maintained at 55.0 ± 0.3 °C, and the latency to first hind paw response (paw lick or shake) was recorded.

### Y-maze test

Spatial working memory and schizophrenia-like behavior were assessed using the Y-maze test, as described previously with minor modifications. The apparatus consisted of three white opaque plastic arms positioned at 120° angles. Each mouse was placed in the center of the maze and allowed to freely explore for 5 min. Arm entries were recorded when all four limbs entered an arm. Spontaneous alternation was calculated as follows: N1/(N2−2), where N1 represents the number of triads containing entries into three different arms, and N2 represents the total number of arm entries.

### Porsolt Forced Swim Test

The Porsolt forced swim test was performed on 9 to 12-wk-old male mice. The apparatus consisted of four plastic cylinders (20 cm height × 10 cm diameter) filled with water (23°C) to a depth of 7.5 cm. Mice were placed individually in the cylinders, and their behavior was recorded for 10 min. Data acquisition and analysis were performed automatically using Image PS software. Distance traveled was quantified by Image OF software.

### Tail suspension test

Depression-like behavior was assessed using the tail suspension test. Mice were suspended 30 cm above the floor with adhesive tape placed 1 cm from the tip of the tail, and their behavior was recorded for 10 min in a visually isolated area. Immobility was automatically quantified using Image TS software.

### Open Field Test and Atomoxetine Administration

Locomotor activity was measured in an open field apparatus (40 × 40 × 30 cm; Accuscan Instruments, Columbus, OH). Nine- to 12-wk-old male mice were placed in the center of the arena, and total distance traveled (in cm), vertical activity (rearing), time spent in the center, stereotyped movement (beam breaks), and fecal boli counts were recorded for 60 min.

For atomoxetine (ATX) experiments, locomotor activity was assessed in a custom-built open field apparatus (40 × 40 × 30 cm). Mice were video-monitored and the distance traveled (in cm) was calculated using ImageJ software. ATX (10 mg/kg, i.p.) or saline was administered 30 min before testing.

### Crawley social interaction test

Social interaction was assessed using a three-chamber apparatus separated by clear Plexiglas walls with small openings between chambers. Mice were habituated to the apparatus for 10 min. One week later, each mouse was habituated again for 10 min before testing. For the object exploration trial, an inanimate object was placed inside a wire cage in one side chamber. For the social interaction trial, the object was replaced with a stranger male C57BL/6N mouse. Time spent near each cage (<4.5 cm) during each 10-min trial was automatically measured using Time CSI2 software.

### Locomotor activity monitoring in home cage

To evaluate circadian locomotor activity, 9 to 12-week-old male mice were monitored in an automated home-cage activity system (O’Hara & Co., Tokyo, Japan). The setup included a standard home cage (29 × 18 × 12 cm) placed beneath an infrared video camera mounted on a 13-cm-high stand. Each mouse was singly housed, and locomotor activity was continuously recorded at 1 frame/s for at least 72 h. Distance travelled was calculated using Image HA software.

### Light/dark transition test

Anxiety-related behavior was assessed using the light/dark transition test (O’Hara & Co., Tokyo, Japan) in 9- to 12-week-old male mice. The apparatus consisted of a box (21 × 42 × 25 cm) divided into equal size of two chambers, lit (390 lux) and dark (2 lux), connected by a small door. Mice were initially placed into the dark chamber and allowed to explore both compartments for 10 min. The number of transitions, time spent in each chamber, latency to first enter the lit chamber, and distance traveled were recorded automatically.

### Inhibitory Avoidance Test

Inhibitory avoidance (IA) test was conducted as described previously ^23^, using an apparatus consisting of a lit chamber (17 × 10 × 21 cm, white walls) connected to a dark chamber (20 × 10 × 21 cm, black walls), via a sliding door. The lit chamber had a white cardboard floor illuminated at 25 klx and the dark chamber contained a metal grid floor illuminated with 940 nm infrared light. Chambers were cleaned with 70 % ethanol between trials.

During training, mice were placed in the lit chamber for 1 min before the door was opened. Once the mouse fully entered the dark chamber, the door was closed, and a foot shock (1.05 mA, 50 Hz, 4 s, 1 ms pulse) was delivered after 10 s. Mice remained in the dark chamber for 1 min after the shock before being returned to their home cage. For memory recall, the same procedure was repeated without shock, and crossover latency (time to enter the dark chamber with all four paws) was recorded.

### Hippocampal slice culture and gene transfection

Hippocampal organotypic slice cultures were prepared from postnatal day 6-9 (P6-9) pups ^24,25^ of I205K knock-in mice or their wildtype littermates of both sexes. Slices were cultured at 35°C on interface membranes (Millipore) and fed with MEM media containing 20% horse serum, 27 mM D-glucose, 6 mM NaHCO_3_, 2 mM CaCl_2_, 2 mM MgSO_4_, 30 mM HEPES, 0.01 % ascorbic acid and 1 μg/ml insulin. pH was adjusted to 7.3 and osmolality to 300-320 mOsm. Slices were biolistically transfected (BioRad) after 5-7 DIV with a plasmid expressing GFP.

### Two-photon microscopy imaging and induction of sLTP in dendritic spines

Time-lapse fluorescence imaging was carried out with a two-photon microscope (FluoView FV1000MPE, Olympus) equipped with two mode-lock femtosecond-pulse Ti:sapphire lasers (MaiTai HP, Sprectra-Physics). Slices were maintained at room temperature (25-27°C) in a continuous perfusion of artificial cerebrospinal fluid (ACSF) containing 119 mM NaCl, 2.5 mM KCl, 3 mM CaCl_2_, 26.2 mM NaHCO_3_, 1 mM NaH_2_PO_4_ and 11 mM glucose, 1 μM tetrodotoxin, 50 μM picrotoxin and 2 mM 4-methoxy-7-nitroindolinyl (MNI)-L-glutamate (Tocris, Bristol, UK) equilibrated with 5% CO_2_/95% O_2_. Imaging was performed at 8-9 DIV in primary or secondary dendrites from the distal part of the main apical dendrite of CA1 pyramidal neurons. sLTP was induced only on thin or small mushroom spines, with a clearly visible head and neck. Two-photon uncaging of MNI-glutamate was performed using 720 nm light (5 mW), with GFP simultaneously excited at 910 nm. Both lasers were aligned daily by imaging and bleaching 0.5 μm fluorescent beads (Polysciences. Inc.). sLTP was induced by 2 ms pulses repeated at 1 Hz for 30 seconds targeted close to the tip of the spine, as previously described ^5^.

At every time-point, a series of 512 × 512-pixel XY-scanned images were taken every 1 μm of depth (Z). The fluorescence intensity of 7-10 of these images were summed to obtain a single Z-stack image. A constant region of interest was outlined around the spine including the spine head and spine neck and the total integrated fluorescence intensity of the green channel after background subtraction was calculated using ImageJ/FIJI (National Institutes of Health) ^19^.

### Human data

Clinical data were obtained from an individual carrying the CAMK2A I205N variant enrolled in a prospective, longitudinal, observational natural history study at the ENCORE CAMK2-GRIN-GRIA Center of Excellence (Erasmus MC, Rotterdam, METC-2021-0099). Following molecular diagnosis by trio-based whole-exome-sequencing, the individual was referred to the CAMK2 Center of Excellence, after the caregivers became aware of the center through the parent organization. Written informed consent was obtained for publication of clinical information and photographs.

### All-atom molecular dynamics simulations

An initial structure of WT is taken from co-crystal structures of CaMKIIα kinase domain in complex with ADP and GluN2B peptides (PDB code: 7UJR) ^13^. Initial structures for other variants (I205K, I205N, F98S and E183V) were gained by residue substitutions using Modeller 10.6 ^26^. The sequences of GluN2Bc peptide used for interaction partners in the simulations are 1295-1310.

All MD simulations were performed with GROMACS2020.2 ^27^ ^28^ and the CHARMM36m force field ^27^. Protein complexes were solvated with TIP3P water ^29^ in periodic simulation boxes with a minimum distance of 1.0 nm between the protein and the box edge. Counter ions were added to neutralize the total system charge. Energy minimization was performed using the steepest descent algorithm for a maximum of 50,000 steps with a step size of 0.01 nm until the maximum force on any atom was less than 1000 kJ/mol/nm. Long-range electrostatic interactions were treated using the particle mesh Ewald (PME) method with a cutoff distance of 1.2 nm ^30^. Van der Waals interactions were smoothly switched off between 1.0 and 1.2 nm. Periodic boundary conditions were applied in all directions. Following energy minimization, the system was equilibrated for 100 ps at constant temperature and volume (NVT ensemble) using the V-rescale thermostat ^31^ and another 100 ps at constant temperature and pressure (NPT ensemble) using the V-rescale thermostat and the C-rescale barostat with a positional restraint applied to protein heavy atoms. Production MD simulations were carried out using the leap-frog integrator with a 2 fs integration time step. Temperature was maintained at 300 K using the V-rescale thermostat, and pressure was maintained at 1 bar using the Parrinello–Rahman barostat with isotropic coupling. All covalent bonds involving hydrogen atoms were constrained using the LINCS algorithm ^32^. Coordinates were saved every 10 ps for subsequent analyses. GPU-accelerated simulations were performed on two hardware architectures: 1) 3× Intel Xeon E E-2468 cpu (8 Cores), Nvidia GeForce RTX 4070 gpu; 2) 1× AMD EPYC 7713 cpu (64 Cores), Nvidia A40 gpu. One 1.5 μs and three 1.0 μs production runs were performed after energy minimization and relaxation simulations.

To calculate RMSD and RMSF during the simulations, trajectories were aligned about the Cα of the kinase domain. RMSD and RMSF analyses were performed using gmx rms and gmx rmsf modules implemented in GROMACS, respectively. RMSD and RMSF of the peptide were calculated with the kinases aligned to determine how the residues of the peptide fluctuated with respect to the kinase. MDAnalysis (version 2.4.3) ^33^ ^34^ was also used to post-process trajectory data and analysis.

### Statistics

Data are represented as mean ± SEM. Student’s t-test or ANOVA with post hoc multiple comparisons was used; p < 0.05 was considered significant.

## Results

### Generation of CaMKIIα I205K Knock-in Mice

To study the functional significance of CaMKII LLPS *in vivo*, we generated knock-in (KI) mice carrying an I205K point mutation at a conserved residue required for binding (Fig. 1A,B) ^13^. AlphaFold modeling predicted that this mutation significantly reduced the inter-chain predicted template modeling (ipTM) score from 0.91 in the wild type (WT) and GluN2B to 0.35 in the I205K mutant (>0.75 is generally considered indicative of a reliable protein–protein interface). Inspection of the predicted model suggested that the I205K mutation may locally perturb the hydrophobic pocket formed by residues F98, I101, V102, and I205^13^, primarily through protrusion of the introduced lysine side chain, without apparent major changes in the overall fold (Fig. 1B).

**Figure 1.**
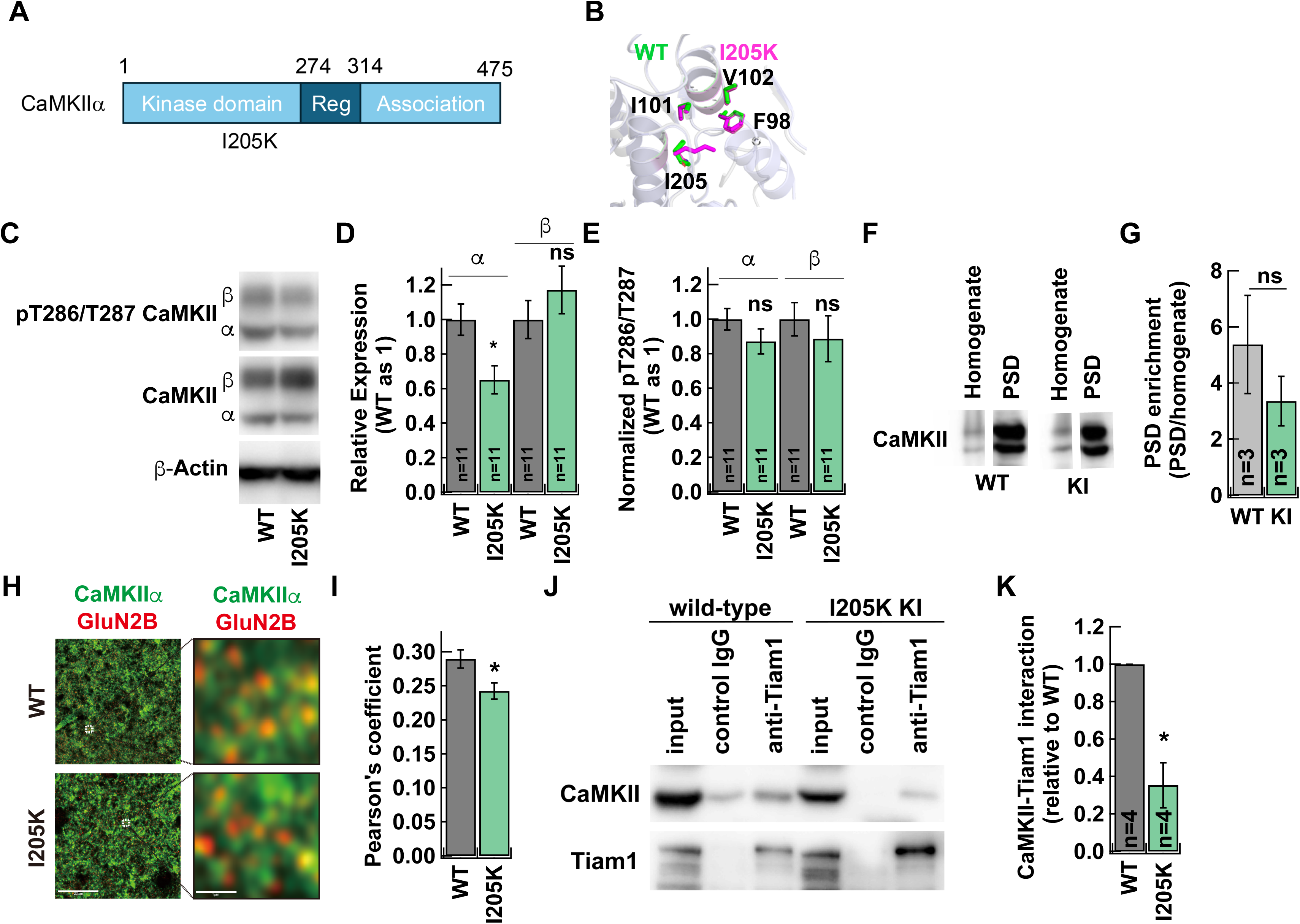
Generation and biochemical characterization of CaMKIIα I205K knock-in mice. **A.** Domain structure of CaMKIIα showing the I205K substitution in the catalytic domain. **B.** Structural models of WT and I205K CaMKIIα hydrophobic pockets in the catalytic domain by AlphaFold3. **C.** Western blot analysis of CaMKII expression in whole-brain homogenate and purified PSD fractions from WT and I205K mice. **D.** Quantification of CaMKII PSD enrichment, calculated as the PSD signal normalized to the homogenate signal (PSD/homogenate). N=3 mice per genotype. **E.** Western blot analysis of CaMKII in brain lysates from WT and I205K mice. Samples were immunoblotted for phosphorylated T286/T287 CaMKII, total CaMKII, and β-actin. **F.** Quantification of CaMKIIα and CaMKIIβ expression normalized to β-actin and shown relative to WT. **G.** Quantification of CaMKII phosphorylation at T286/T287 normalized to total CaMKII. **H.** Representative confocal images of CaMKIIα (green) and GluN2B (red) immunostaining in the hippocampal CA1 region. **I.** Quantification of CaMKIIα and GluN2B co-localization using Pearson’s coefficient (5 images per animal, n=2 mice). Scale bars: left, 20 µm; right, 10 µm. **J.** Co-immunoprecipitation from adult mouse brain lysate using an anti-Tiam1 antibody, following by immunoblotting for CaMKII and Tiam1. **K.** Quantification of co-immunoprecipitated CaMKII normalized to WT (n=4, mice per genotype). Data are shown as mean ± SEM. ns, not significant, *, p<0.05, in a two-tailed unpaired t-test, compared to WT.

In brain lysates, total CaMKIIα levels were significantly reduced in I205K mice (p=0.019), whereas CaMKIIα expression was comparable to WT (p=0.143; Fig. 1C,D). Autophosphorylation of CaMKIIα at T286 did not differ significantly between genotypes (p=0.323; Fig. 1C,E). In purified postsynaptic density (PSD) fractions, CaMKIIα from I205K mice showed a modest reduction relative to WT, although this difference was not statistically significant (p=0.069; Fig. 1F,G). Immunostaining of the hippocampal CA1 region showed decreased co-localization of CaMKIIα with GluN2B in KI mice compared with WT (p=0.0033; Fig. 1H,I). Together, these results indicate that the I205K mutation weakens CaMKII-GluN2B association and synaptic enrichment while leaving CaMKII autophosphorylation largely intact^35^.

To determine whether this mutation disrupts interactions between CaMKIIα and binding partners other than GluN2B, we performed co-immunoprecipitation of CaMKII with Tiam1 from brain lysates of KI and WT mice. We observed a marked reduction of CaMKIIα in the Tiam1 precipitate from KI mice compared with WT (p=0.0036; Fig. 1J,K), indicating reduced CaMKIIα-Tiam1 interaction in the mutant. These results suggest that the I205K mutation primarily impairs protein localization and binding interactions rather than kinase activation itself^35^.

### Spine morphology and glutamate receptor distribution in CaMKIIα I205K knock-in mice

We next asked whether the I205K mutation affects synaptic structure. Golgi-Cox staining was used to visualize pyramidal neurons in the CA1 region. Quantitative analysis revealed no significant differences in spine density, head diameter, or spine neck length between KI and WT mice (p=0.242, p=0.3621, p=0.054, respectively; Fig. S1), indicating that the I205K mutation does not substantially affect neuronal maturation or dendritic spine morphology under basal conditions.

CaMKII-mediated LLPS has been proposed to induce developmental and activity-dependent segregation of AMPA receptor (AMPAR) and NMDA receptor (NMDAR) nanodomains at synapses *in vitro* ^16,36^. We therefore investigated whether I205K mice exhibit altered synaptic receptor segregation *in vivo* using Sodium Dodecyl Sulfate-Digested Freeze-Fracture Replica Labeling (SDS-FRL). We visualized the distribution of AMPARs and NMDARs with immunogold particles of different sizes. PSD area was demarcated by the presence of intramembrane particles (Fig. 2A) ^37,38^. The average PSD area per spine and the density of each receptor were comparable between genotypes (PSD area: p=0.623; AMPAR: p=0.257; NMDAR: p=0.199; Fig. S2). We next analyzed nearest-neighbor distances to assess the spatial organization and clustering of receptors within the PSD. Increased inter-NMDAR distances were observed in dendritic spines of KI mice (p=0.012; Fig. 2B right), whereas AMPAR nearest-neighbor distances were unchanged (p=0.145; Fig. 2B left). The larger NMDAR distances align with the modest reduction in NMDAR density observed in KI spines (Fig. S2), even though this density change did not reach statistical significance. We also compared the nearest-neighbor distances between AMPARs and NMDARs, which were significantly greater than those in the simulated random distribution in both WT (p<0.0001) and KI mice (p<0.0001; Fig. 2C). These findings suggest that AMPARs and NMDARs were spatially segregated within PSDs in both genotypes, consistent with non-random receptor organization and previous observations using super-resolution microscopy ^16^.

**Figure 2.**
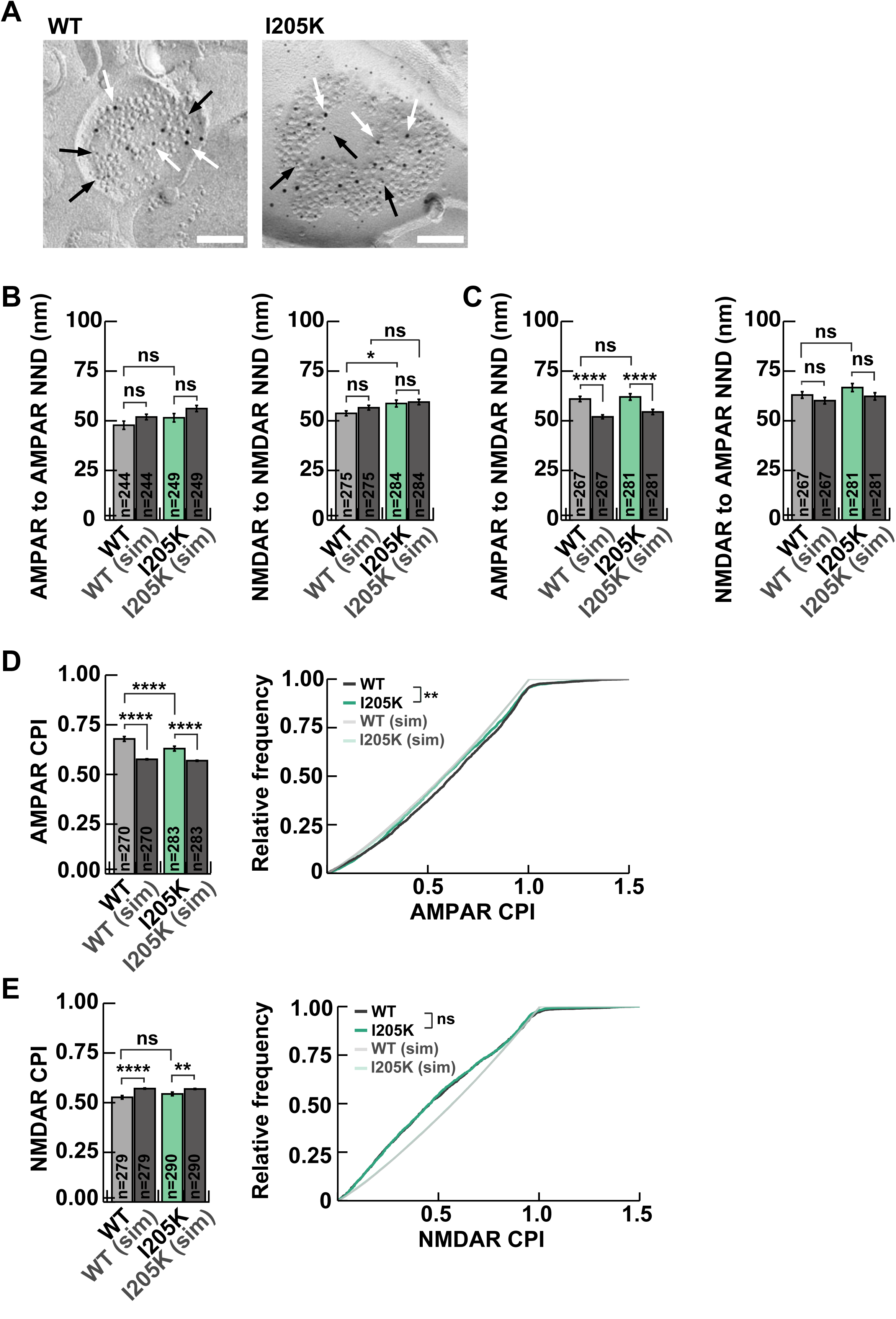
Distribution of AMPA and NMDA receptors within the PSD analyzed by SDS-FRL. **A.** Representative SDS-FRL images of WT and I205K synapses showing immunogold labeling of AMPARs (5-nm gold; black arrows) and NMDARs (10-nm gold; white arrows). Scale bar, 100 nm. **B.** Nearest-neighbor distance analysis showing clustering of NMDARs within synapses in WT mice compared to I205K mice (WT, n=275; I205K, n=284 synapses), whereas AMPARs did not significantly differ from simulated values (WT, n=283; I205K, n=292 synapses). **C.** Nearest-neighbor distances between AMPARs and NMDARs (heterotypic pairs) exceeded simulated values in both WT and I205K mice, suggesting spatial segregation between the two receptor types (WT, n=267; I205K, n=281 synapses). **D.** The Center–Periphery Index (CPI) for AMPARs was greater than the simulated value in both WT and I205K mice. The CPI frequency distribution for AMPARs was right-shifted in WT mice relative to I205K mice, indicating more peripheral positioning of AMPARs in WT mice (WT, n=270; I205K, n=283 synapses). **E.** The CPI for NMDARs was smaller than simulated values in both WT and I205K mice. CPI distributions for NMDARs were left-shifted in WT and I205K mice relative to simulated values, indicating central localization of NMDARs (WT, n=279; I205K, n=290 synapses). Data are shown as mean ± SEM. Statistical significance was determined using a two-tailed unpaired t-test, two-way ANOVA with post-hoc multiple comparisons and Kolmogorov-Smirnov test (*p < 0.05, **p < 0.01, ***p < 0.001, ****p < 0.0001, ns = not significant.).

For further analysis, we examined the relative radial position of each receptor within PSDs using the Center-Periphery Index (CPI) ^18^. The mean CPI for AMPARs was significantly larger than that of the simulated random distribution in both WT (p<0.0001) and KI (p<0.0001) mice, and was significantly higher in WT than in KI mice (p<0.0001; Fig. 2D left). This indicates that AMPARs are preferentially located toward the periphery of the PSD and that this peripheral enrichment was reduced in KI mice. In contrast, the mean CPI for NMDARs was significantly smaller than that of the simulated values (WT: p<0.0001; KI: p=0.0059), indicating preferential localization toward the center (Fig. 2E left). No significant difference was observed between the genotypes (p=0.068). The CPI frequency distribution for AMPARs was, consistently, left-shifted in KI mice compared to WT mice (p=0.0052; Fig. 2D, right) while the distribution of NMDARs was nearly identical between WT and KI mice (p=0.859; Fig. 2E, right).

In summary, NMDARs were located preferentially toward the center of the PSD in both genotypes, whereas AMPARs were positioned toward the periphery in WT mice. This peripheral enrichment of AMPARs was reduced in KI mice, resulting in weaker radial separation between peripheral AMPARs and the centrally positioned NMDARs. Thus, while overall AMPAR-NMDAR segregation is preserved *in vivo*, the I205K mutation diminishes the peripheral positioning of AMPARs relative to a stable NMDAR scaffold, indicating that CaMKII-GluN2B interaction plays a role in refining the postsynaptic nano-architecture ^36^.

### Impaired structural LTP in I205K knock-in mice

We next assessed the impact of this mutation on activity-dependent modification of the synapse by examining sLTP. GFP was biolistically expressed in CA1 pyramidal neurons of hippocampal slices, and spine enlargement was induced by two-photon glutamate uncaging. In slices from WT mice, stimulated spines exhibited persistent volume increases, whereas spines from KI slices showed only a transient enlargement that decayed to baseline within 5 min (Fig. 3A,B). Group data revealed that both the transient (1-3 min) and sustained (21-29 min) phases of sLTP were significantly reduced in KI neurons compared with WT (p=0.024; p=0.0044, respectively; Fig. 3C), demonstrating that the I205K mutation impairs activity-dependent structural plasticity. These findings are consistent with previous reports showing that the I205K mutant fails to rescue LTP deficits in CaMKIIα/β double-knockout neurons ^39^ and further support the model that CaMKII-GluN2B interaction mediated by the hydrophobic interaction is critical for effective synaptic potentiation ^12,40^.

**Figure 3.**
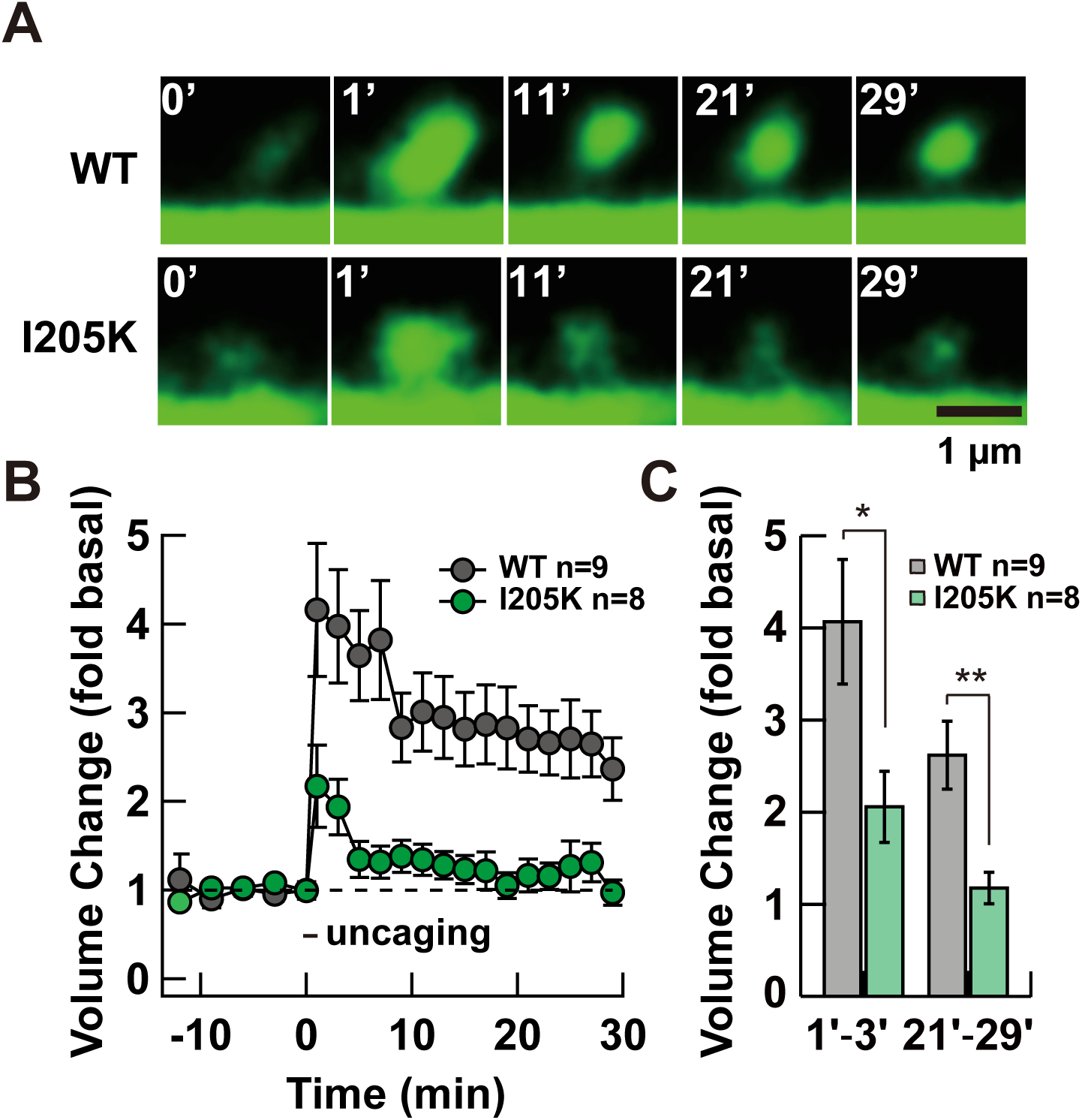
Impaired structural plasticity in hippocampal neurons of CaMKIIα I205K knock-in mice. **A.** Representative two-photon images showing sLTP in hippocampal pyramidal neurons expressing GFP in organotypic slice cultures from WT (top; n=9 spines) and I205K (bottom; n=8 spines) mice. Scale bar, 1 µm. **B.** Time course of spine volume change during sLTP in WT and I205K neurons. **C.** Quantification of the transient phase (1-3 min) and sustained phase (21-29 min) of sLTP. Data are shown as mean ± SEM. *, p<0.05, **, p<0.01 in a two-tailed unpaired t-test, compared to WT.

### Behavioral battery tests

We next asked whether these synaptic deficits translate into alterations in physiology, motor function and cognitive behaviors. We therefore performed a comprehensive behavioral test battery on WT and CaMKII I205K KI mice. KI mice were viable but significantly leaner than WT littermates (p<0.001; Fig. 4A). KI mice also exhibited a lower rectal temperature (p=0.04; Fig. 4B). In response to thermonoxious stimuli, as evaluated by the hot plate test (p<0.001; Fig. 4C), KI mice showed a shorter reaction latency, indicating heightened sensitivity. KI mice also displayed reduced motor performance, as assessed by grip strength (p<0.001; Fig. 4D) and the wire hang test (p<0.001; Fig. 4E).

**Figure 4.**
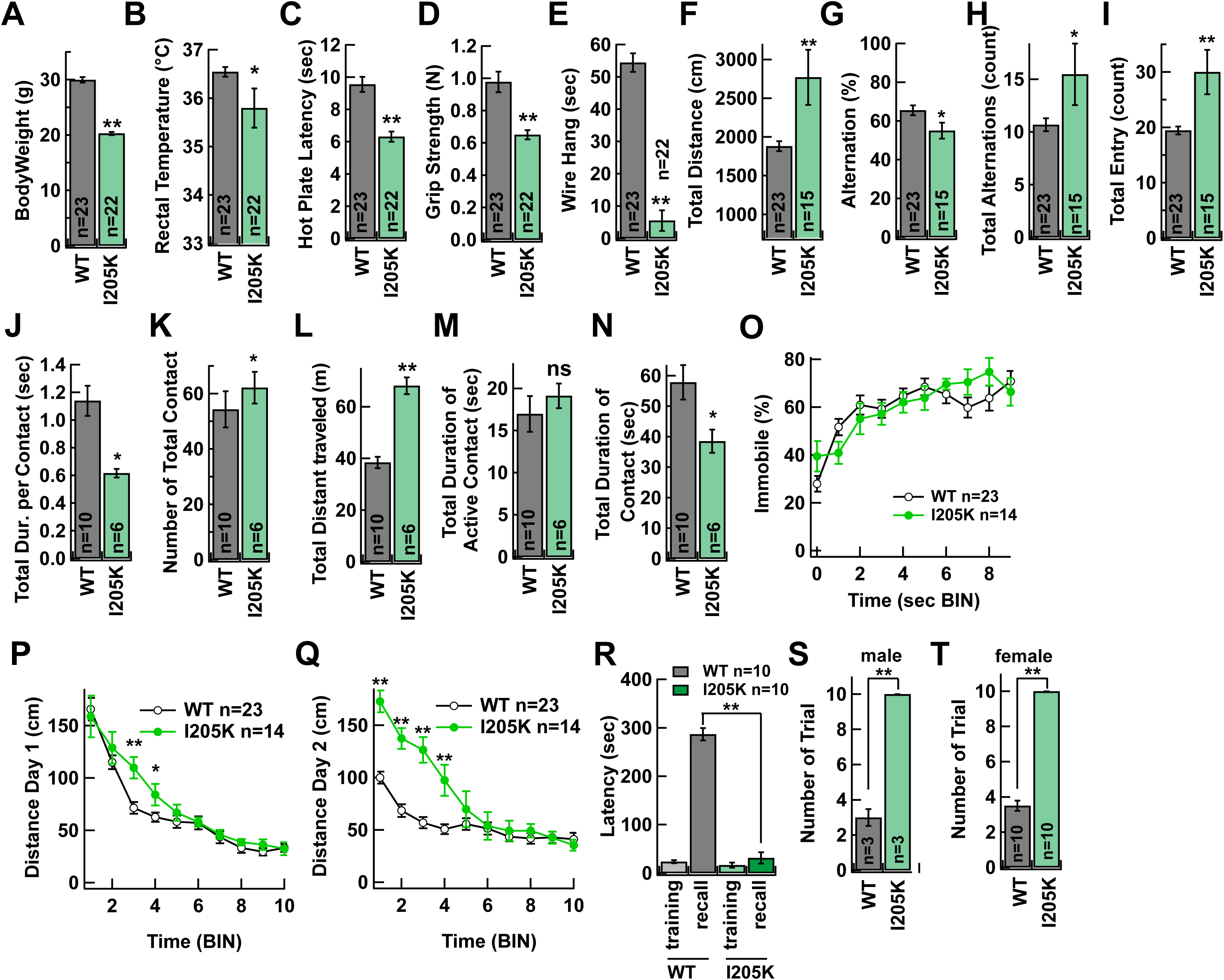
Behavioral characterization of CaMKIIα I205K knock-in mice A-E. General health and motor function: (A) Body weight (g). (B) Rectal temperature (°C). (C) Hot plate latency (sec). (D) Grip strength in Newton (N). (E) Wire-hang test (sec) (WT, n=23; I205K, n=22). **F-I**. Y-maze test (WT, n=23; I205K, n=15). (F) total travel distance (cm). (G) Alteration to novel arm (%). (H) Total alteration number (count). (I) Toral entry number (count). **J-N**. Social interaction test (WT, n=10; I205K, n=6). (J) total duration per contact (sec). (K) Number of total contact (count). (L) Total distant traveled (m). (M) Total duration of active contact (sec). (N) Total duration of contact (sec). **O**. Tail-suspension test (WT, n=23; I205K, n=14). Immobile time during recording (%). **P-Q.** Forced swim test (WT, n=23; I205K, n=14). (P) Swimming distance at Day 1 (cm). (Q) Swimming distance at Day 2 (cm). **R**. Inhibitory avoidance task showing crossover latency in training and recall sessions. Crossover latency for training and recall trials were measured. Cutoff latency set at 300 sec. (WT, n=10; I205K, n=10) **S**-**T**. Number of trainings trials required to cross to the dark chamber. (S) Male mice of WT or I205K. Cutoff trial set at 10 training. (WT, n=3; I205K, n=3). (T) Female mice of WT or I205K. (WT, n=10; I205K, n=10) Data are shown as mean ± SEM. *p < 0.05, **p < 0.01, ns = not significant. Two-tailed unpaired t-test, compared to WT.

Spatial working memory was investigated using the Y-maze (Fig. 4F-I). While WT mice preferentially explored the novel arm rather than the familiar one, KI mice showed no preference, performing close to chance level. In addition, KI mice exhibited increased locomotor activity, reflected by greater total distance travelled (p=0.002; Fig. 4F), total alternation (p=0.031; Fig. 4H), and total arm entries (p=0.002; Fig. 4I).

In the social interaction test (Fig. 4J-N), KI mice showed shorter contact duration (p=0.029; Fig. 4N) but a greater number of contacts compared with WT mice (p=0.029; Fig. 4K).

Depression-like behavior and behavioral despair were evaluated using the tail suspension test (TST; Fig. 4O) and the forced swim test (FST; Fig. 4P,Q). Immobility time was not significantly different between genotypes, indicating normal depression- and despair-related behaviors. However, in the FST, KI mice took slightly longer to exhibit immobility on day 1 (Fig. 4P). On day 2, while WT mice adapted to the test and reached immobility faster than on day 1, KI mice showed similar immobility curves across both days (Fig. 4Q), suggesting impaired learning or memory of the test environment.

Finally, associative learning was examined using the inhibitory avoidance (IA) test (Fig. 4R). WT mice remembered the aversive experience as indicated by prolonged hesitation to enter the dark chamber, whereas KI mice entered the dark compartment without latency (recall latency WT vs KI, p<0.001; Fig. 4R). We also conducted repeated IA training with reduced shock intensity to determine whether learning could be achieved through multiple exposures. WT mice acquired the aversive memory within three sessions, while I205K mice failed to show avoidance behavior even after more than ten trials in both males (p<0.001; Fig. 4S) and females (p<0.001; Fig. 4T). These findings indicate that the I205K mutation leads to severe deficits in aversive learning and memory, distinct from the phenotype observed in CaMKII T286A knock-in mice, which are capable of learning with repeated training ^41^.

### CaMKIIα I205K knock-in mice show ADHD-like hyperactive behavior

Having tested these, we became aware that the interpretation of these behavioral results could be compromised by the overall hyperactivity of the KI animals. The hyperactivity was obvious in the open field (Fig. 5A,B) and the home cage (Fig. 5C) upon tracking mouse trajectories. We found that the mice continuously moved along the wall, which explains the reduced time spent in the center (Fig. 5B). Additionally, when we attempted to test the elevated plus maze, KI mice jumped off the maze and were unable to perform the task (not shown). This hyperactivity may underlie their lower body mass, including reduced adipose tissue, which could also contribute to the reduced body temperature.

**Figure 5.**
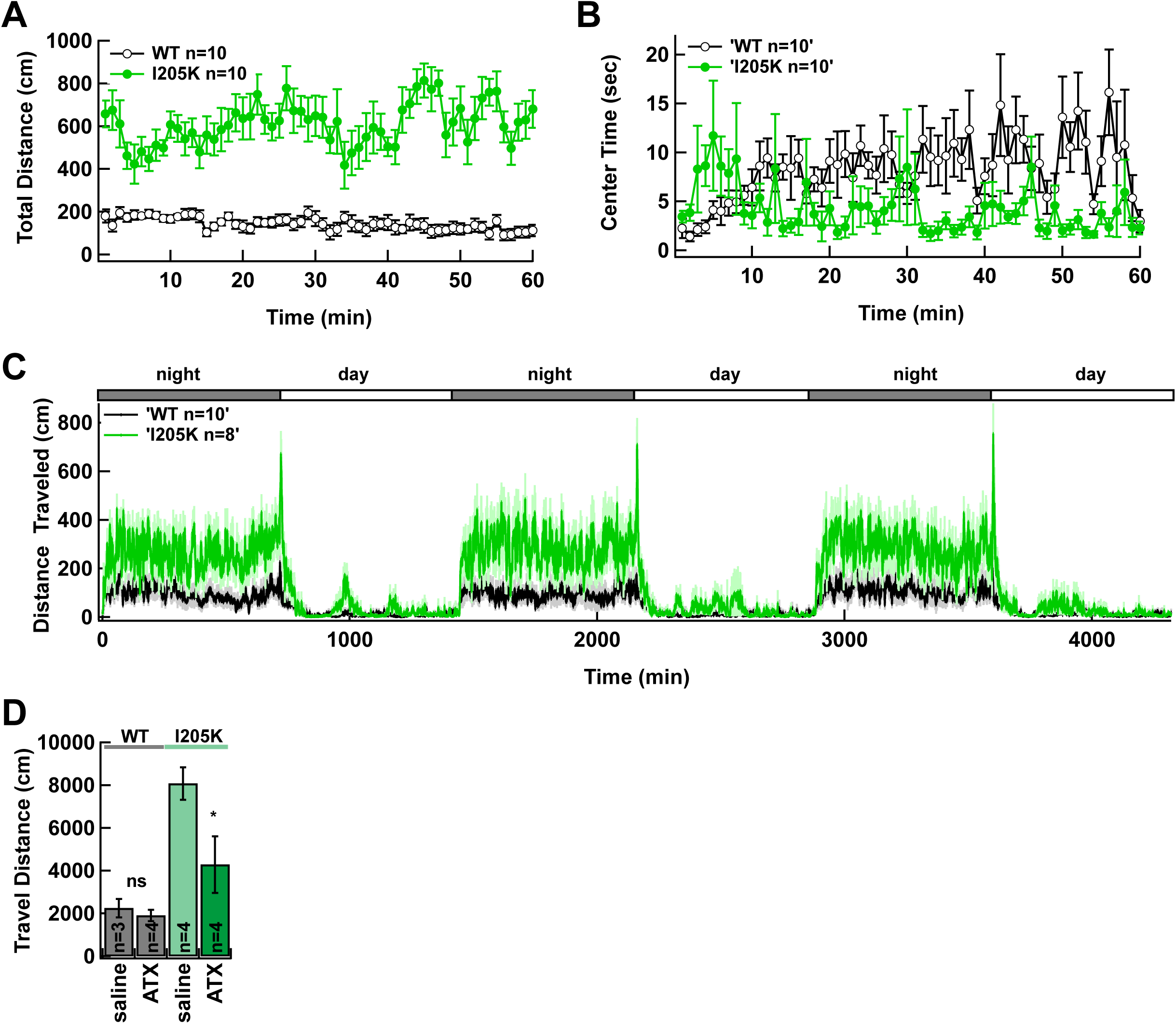
Hyperactivity and learning impairment in CaMKIIα I205K knock-in mice. **A-B**. Locomotor activity levels were tested in open field test (WT, n=10; I205K, n=10). (A) total distance (cm). (B) Time spent in center of the arena (sec). **C**. Home-cage activity monitored for 72 hours showing total distance traveled (cm) (WT, n=10; I205K, n=8). **D**. Effect of atomoxetine (ATX; 10 mg/kg, intraperitoneal, 30 min before test)) on locomotor activity (WT: saline n=3, ATX n=4; I205K: saline n=4, ATX n=4). Data are shown as mean ± SEM. *, p<0.05, in a two-tailed unpaired t-test, compared to saline vs ATX.

These findings reminded us of the persistent hyperactivity seen in attention-deficit/hyperactivity disorder (ADHD). To test whether the hyperactive phenotype shares any common mechanisms with ADHD, we examined the effect of the norepinephrine-selective reuptake inhibitor, atomoxetine (ATX), an FDA-approved treatment for ADHD (Fig. 5D). ATX treatment did not alter WT behavior (p=0.291) but significantly reduced hyperactivity in KI mice in the open field (p=0.038), supporting the idea that the I205K mutation may model ADHD-like behavioral abnormalities.

### Disruption of CaMKII Phase Separation in Human with CaMKII Variants

Because CaMKIIα I205K knock-in mice exhibited neurodevelopmental disorder-like behavioral abnormalities, we hypothesized that variants in the hydrophobic pocket define a potential neurodevelopmental disorder-risk domain within CaMKIIα. The hydrophobic pocket of CaMKIIα interacting with GluN2B are comprised of four key residues, F98, I101, V102, and I205 (F99, I102, V103, and I206 in other subunits) ^13^. Analysis of publicly available whole-exome sequencing datasets identified several *de novo* missense variants within this hydrophobic pocket in patients with neurodevelopmental disorders. The F98S variant has been reported in individuals with intellectual disability ^42^. From the PRISM functional genomic consortium, we identified an additional novel human neurodevelopmental disorder-associated variant in this region, I205N, in a 13-year-old male, who was diagnosed at 6 years of age (Table S1). Developmental milestones were globally delayed, prompting initial evaluation at age 2. At 13 years, he exhibited mild intellectual disability, with speech intelligibility mildly impacted by dyspraxic and dysarthric features. Gross motor function was generally adequate in daily activities, formal testing, however, demonstrated motor delay, with more pronounced fine motor difficulties. He was diagnosed with ADHD, with accompanying difficulties in stimulus processing, consistent with findings in the mouse model. Socially, he demonstrated preserved attunement to others and appropriate reciprocal interaction. He had no history of seizures, although an EEG demonstrated focal epileptiform activity.

To explore whether such neurodevelopmental disorder-associated variants share functional similarities with the I205K mutant, particularly in their ability to undergo liquid-liquid phase separation (LLPS) with GluN2B, we analyzed F98S and I205N. Additionally, we also examined E183V, which is associated with autism spectrum disorder but without a clearly proposed structural mechanism ^43^. To assess LLPS formation, purified GFP-tagged CaMKIIα WT or variants were mixed with the oligomeric GluN2B cytoplasmic tail and calmodulin in the presence of Ca^2+^ and ATP/Mg^2+^. Confocal imaging revealed that WT formed clear, spherical condensates (∼5 µm in diameter), whereas I205K failed to form condensates under the same conditions, consistent with previous findings ^16^. Likewise, all neurodevelopmental disorder-associated variants, including F98S, E183V, and I205N, failed to form condensates with GluN2B (Fig. 6A), indicating that the impairment of phase separation is a common abnormality caused by the human variants leading to neurodevelopmental disorders. We next assessed these interactions by co-immunoprecipitation from HEK293T cells co-expressing FLAG-tagged CaMKII WT or variants and the cytoplasmic tail of GluN2B (GluN2Bc). Following immunoprecipitation with an anti-FLAG antibody, co-precipitated proteins were detected with a GluN2B antibody. WT CaMKIIα robustly co-precipitated with GluN2B, whereas I205K and other variants showed markedly reduced interactions (Fig. 6B,C). Notably, E183V expression was consistently lower than that of WT or other variants suggestive of impairment in protein expression or stability ^43^.

**Figure 6.**
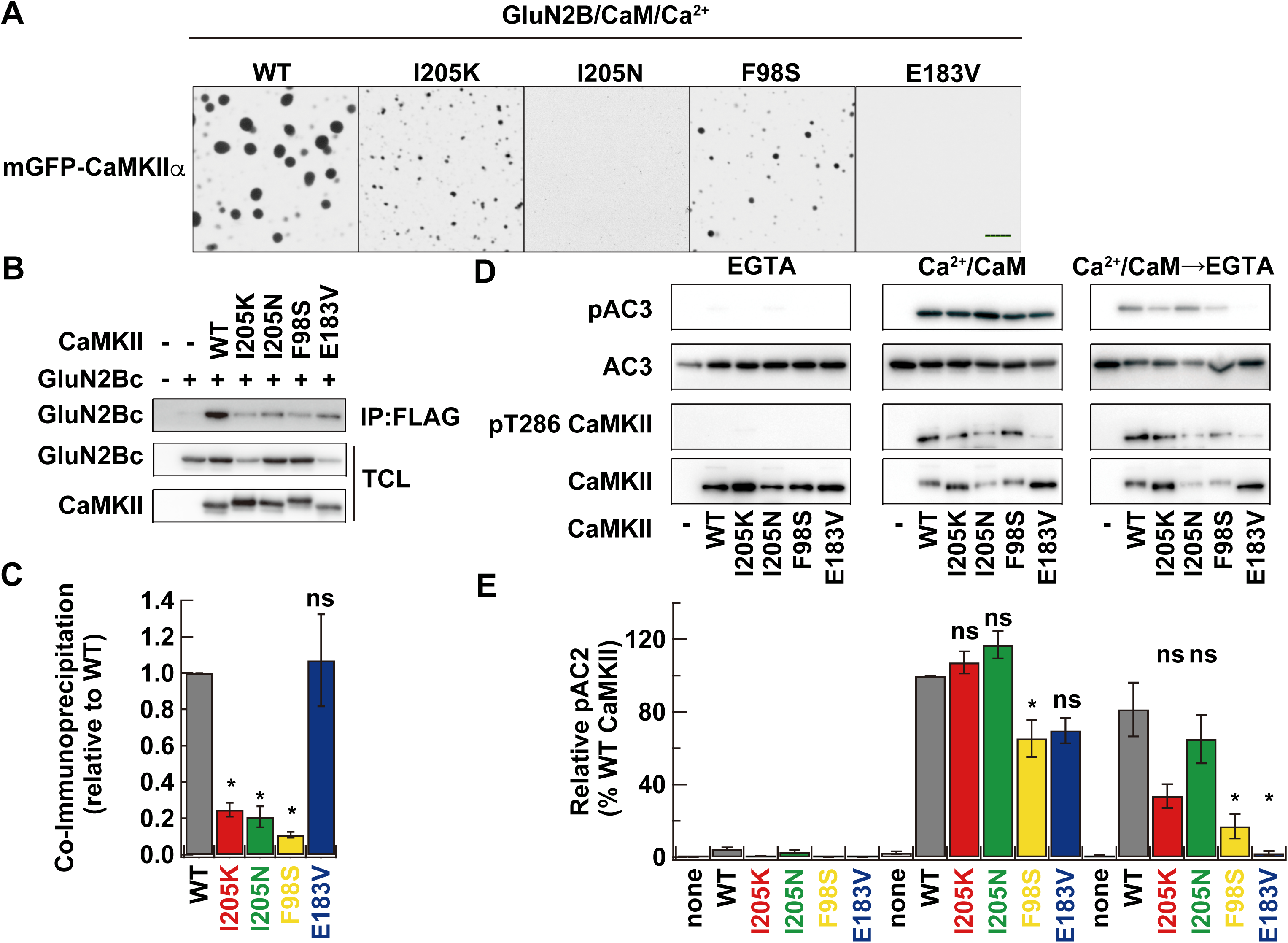
Impaired LLPS and interactions in I205K and neurodevelopmental disorder-associated variants. **A.** Confocal images showing phase separation of GFP-tagged CaMKIIα WT, I205K and neurodevelopmental disorder-associated variants (I205N, F98S, E183V) in the presence of Mg^2+^/ATP, Ca^2+^/CaM, and oligomeric GluN2B. Scale bar, 5 µm. **B.** Co-immunoprecipitation analysis of the interaction between CaMKIIα and GluN2B. HEK293T cell lysates expressing FLAG-tagged CaMKIIα WT or the indicated variants (I205K, I205N, F98S, and E183V), together with GFP-tagged GluN2B cytoplasmic tail (aa 1120-1482), were immunoprecipitated with anti-FLAG antibody and immunoblotted for CaMKIIα and GluN2B. IP, immunoprecipitate; TCL, total cell lysate. **C.** Quantification of co-immunoprecipitated GluN2B normalized to WT level (n=4). Data are presented as mean ± SEM. ns, not significant, *, p<0.05, in a two-tailed unpaired t-test, compared to WT. **D.** In vitro kinase assay using the AC2 peptide substrate. Purified AC2 peptide was phosphorylated by GFP-CaMKII in the absence (EGTA) or presence of Ca^2+^/CaM. Autonomous activity was assessed after preincubation of CaMKII with Mg^2+^/ATP and Ca^2+^/CaM, followed by addition of EGTA. **E.** Quantification of kinase activity normalized to WT (Ca^2+^/CaM condition = 1; n=6 for WT, I205K, and I205N; n=3 for F98S and E183V). Data are shown as mean ± SEM. *, p<0.05, in a two-tailed unpaired t-test, compared to WT.

We examined the enzymatic properties of CaMKIIα WT, I205K and the neurodevelopmental disorder-associated variants using the AC2 peptide as a substrate ^44^. Under Ca^2+^-free conditions, no phosphorylation of AC2 was detected in any samples (Fig. 6D,E). In the presence of Ca^2+^, the mutants exhibited kinase activities comparable to WT, except for F98S, which showed reduced activity. Chelation of Ca^2+^ by EGTA revealed that I205K and I205N retained WT-like autonomous activity, whereas F98S and E183V displayed significantly lower activity (Fig. 6E). Autophosphorylation at T286 was largely comparable between WT and the variants, though E183V exhibited a noticeable reduction under both Ca^2+^ and EGTA conditions ^43^.

### Molecular dynamics simulations reveal distinct mechanisms destabilizing the CaMKII–GluN2B interaction

Both I205 and F98 constitute a hydrophobic patch important for binding with GluN2B, and it is therefore reasonable that mutations in these residues impair LLPS with GluN2B. However, the mechanism by which E183V impairs LLPS remained unclear, given that E183 is not directly involved in the interaction with GluN2B ^13^. To understand how these CaMKII variants disrupt LLPS, we carried out AlphaFold3 prediction. The I205K and F98S variants showed significant reductions in ipTM score compared to WT (WT: 0.91, I205K: 0.35, and F98S: 0.71), whereas the I205N and E183V variants did not show appreciable reductions (I205N: 0.92 and E183V: 0.91), which failed to explain the observed impairment in LLPS. On the other hand, on the AlphaMissense database ^45^, they are all highly pathogenic (>0.99). As AlphaFold3 prediction primarily evaluates the static structural compatibility of protein-protein interfaces and does not fully capture the dynamic stability of intermolecular interactions, we performed molecular dynamics (MD) simulations to further investigate how these mutations affect the persistence and stability of the CaMKII–GluN2B interaction over time. We performed all-atom MD simulations of the CaMKII kinase domain (residues 1–274), WT or variants, in complex with the GluN2B peptide (residues 1295–1310) and ADP.

In the WT complex, the interaction interface remained highly stable throughout the simulation (Fig. 7, Fig. S3). The canonical interaction motifs, including the +1 hydrophobic patch, −2 hydrogen bond, −3 salt bridge, −5 hydrophobic pocket, and −8 salt bridge, were largely immobile over trajectories, consistent with a stable GluN2B-bound conformation (Fig. 7B-D). In contrast, the I205K mutation induced marked destabilization of the interaction interface (Fig. 7B-D, Fig. S3). MD trajectories revealed progressive displacement of the GluN2B peptide accompanied by dissociation of the +1 hydrophobic interaction and disruption of the −2 hydrogen bond, −3 salt bridge, and −8 salt bridge interactions (Fig. 7 and Fig. S3). The hydrophobic pocket surrounding residue 205 underwent substantial rearrangement during the simulation, resulting in broad loss of intermolecular contacts in contact map analyses (Fig. S3A). Notably, these structural changes emerged dynamically during the simulation despite preservation of the overall kinase fold, suggesting that the mutation primarily impairs interaction stability rather than causing global protein unfolding.

**Figure 7.**
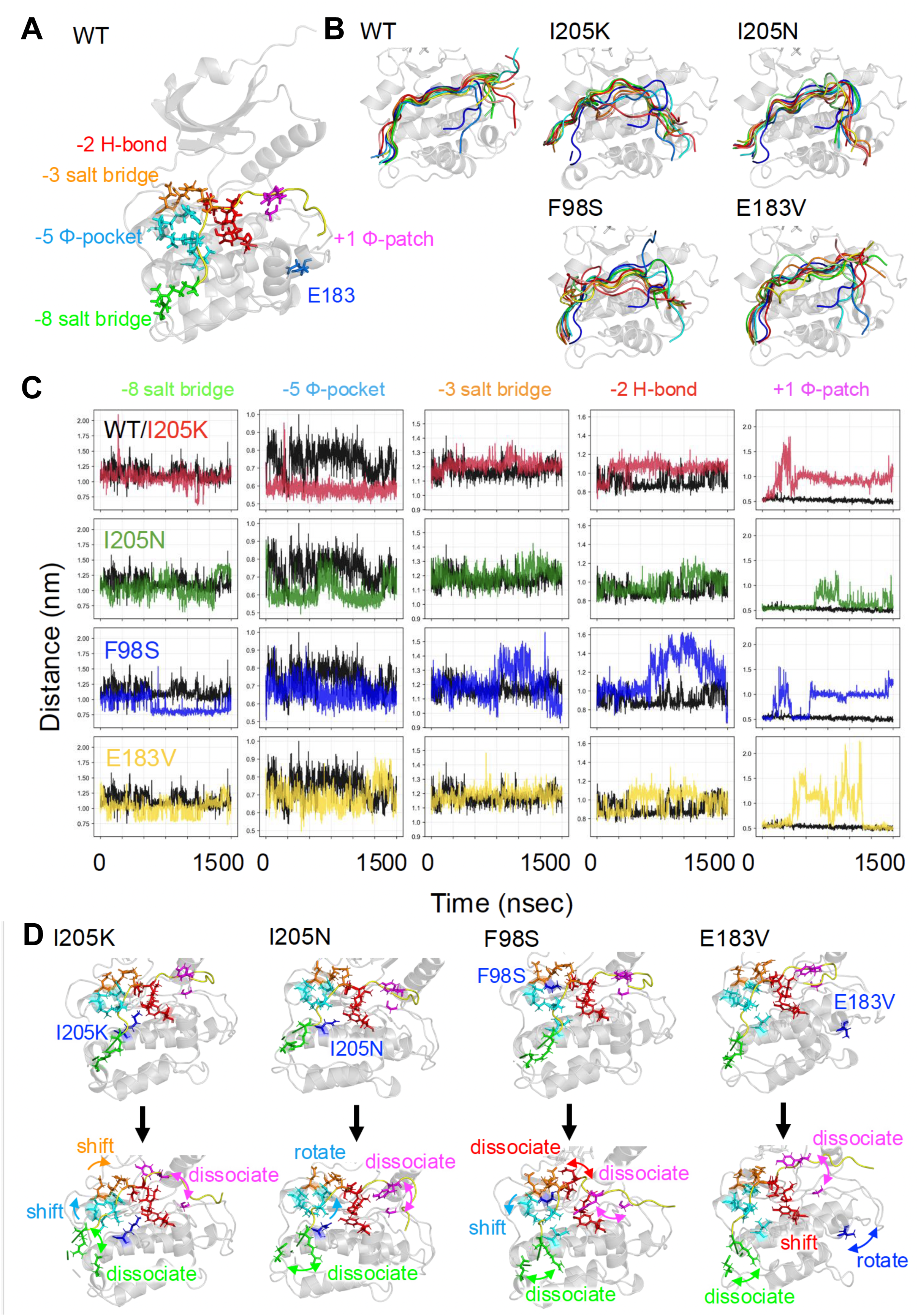
Molecular dynamics simulations of the CaMKII–GluN2B interaction. **B.** Structure of the CaMKII kinase domain (gray) bound to the GluN2Bc peptide. Residue numbering of the peptide is defined relative to Ser1303 (position 0). A salt bridge in -8 (green), a hydrophobic pocket (Φ-pocket) in -5 (cyan), a salt bridge in -3 (orange), a hydrogen bond in -2 (red) and hydrophobic patch (Φ-patch) in +1 (magenta) are shown. Residue 183 is located in the distant area from interactions site (blue). **C.** Dynamics of GluN2Bc peptides in WT and variants. The colors correspond to the timeline, ranging from initial (blue) to final (red). **D.** Distances between the five interaction site residues and their corresponding GluN2Bc residues in representative trajectories for WT (black), I205K (red), I205N (green), F98S (blue), and E183V (yellow). The salt bridge at -8 indicates the distance between E236 and R-8; the hydrophobic pocket at -5 indicates the distance between V102 and L-5; the salt bridge at -3 indicates the distance between E96 and R-3; the hydrogen bond at -2 indicates the distance between E139 and Q-2; and the hydrophobic patch at +1 indicates the distance between G175 and Y+1. **E.** Changes observed before (top) and after (bottom) major structure shift occurred in each variant.

The I205N variant exhibited a distinct but related phenotype. Although AlphaFold3 predicted a largely preserved interface for I205N, MD simulations demonstrated increased conformational fluctuations of GluN2B peptide within the hydrophobic pocket and intermittent dissociation of the +1 hydrophobic interaction (Fig. 7B). In particular, rotation and rearrangement of hydrophobic side chains near the −5 interaction site destabilized peptide binding without inducing the severe interface collapse observed in I205K (Fig. S3). These findings may explain why I205N retains relatively preserved predicted structural compatibility while still impairing LLPS experimentally.

The F98S variant exhibited a distinct mode of interface destabilization. Unlike I205K, which caused marked displacement of the GluN2B peptide, F98S largely preserved the overall binding geometry but weakened hydrophobic packing within the interaction interface. Replacement of the bulky hydrophobic phenylalanine residue with serine reduced the stability and persistence of local intermolecular contacts, particularly around the +1 hydrophobic, -2 hydrogen bond and -3 salt bridge interaction region (Fig. 7D, Fig. S3). These findings suggest that subtle weakening of hydrophobic interaction networks, even without large-scale structural rearrangement, is sufficient to impair the multivalent interactions required for CaMKII-mediated LLPS.

Interestingly, the E183V variant, which is spatially separated from the GluN2B interaction interface, also exhibited altered interaction dynamics in MD simulations. Rather than directly disrupting the canonical GluN2B binding site, E183V induced displacement of surrounding loop structures, resulting in secondary destabilization of the hydrophobic interaction network. Structural analysis revealed that E183 forms a stable intramolecular salt bridge with R259, with the carboxylate oxygens of E183 positioned within salt-bridge distance of the guanidinium nitrogens of R259 (2.65 Å and 2.82 Å. Fig.S3B). This E183–R259 interaction appears to clamp the neighboring loop region adjacent to the GluN2B-binding surface. Consistent with this structural arrangement, E183V mutation displaced the local loop outward during the simulation, thereby disrupting the geometry of the adjacent GluN2B-contacting surface and promoting dissociation of the hydrophobic interface involving the CaMKII G175-region and the GluN2B (Fig. 7C). These results suggest that E183 contributes to GluN2B binding indirectly by stabilizing the local CaMKII conformation through an E183–R259 electrostatic network.

Together, these simulations suggest that disease-associated CaMKII variants disrupt LLPS not only through gross loss of binding affinity, but also by perturbing the dynamic stability of multivalent interaction interfaces required for persistent molecular assembly.

## Discussion

This present study demonstrates that CaMKIIα phase separation is a key mechanism for organizing postsynaptic signaling and enabling activity-dependent plasticity *in vivo*. The I205K mutation reduced CaMKIIα enrichment at the postsynaptic density and its association with GluN2B without altering autophosphorylation, and was further accompanied by refined nanoscale organization of postsynaptic receptors, impaired structural LTP, deficits in aversive learning, and hyperactivity that was reversible with atomoxetine. Human neurodevelopmental disorder–associated variants within the CaMKIIα hydrophobic pocket similarly disrupted phase separation while largely preserving kinase activity. Together, these findings establish CaMKII LLPS as a mechanistic basis for postsynaptic organization, synaptic plasticity and behavior, and implicate its disruption in neurodevelopmental disease.

### Activity-dependent synaptic dysfunction despite preserved basal architecture

The lack of major changes in spine morphology (Fig. S1), or overall synaptic ultrastructure (Fig. S2) suggests that the I205K mutation largely preserves basal synaptic architecture. In contrast, the nearly abolished structural LTP (Fig. 3), together with previously reported impairment of functional LTP ^39^, indicates a preferential defect in activity-dependent synaptic remodeling. This dissociation is consistent with the prominent role of CaMKIIα in mature excitatory neurons, where it may primarily support activity-dependent postsynaptic reorganization rather than basal synapse formation or maintenance. The observed learning deficits and hyperactivity (Fig. 4,5) may therefore arise from disrupted plasticity across hippocampal and extra-hippocampal circuits, rather than from gross abnormalities in basal synaptic structure. Because neuronal activity also contributes to synapse maturation and circuit refinement ^46^, subtle developmental contributions cannot be excluded. Further studies using region-specific analyses of activity-dependent structural remodeling and circuit dynamics will be important to clarify these possibilities.

### Synaptic biomolecule condensates formation and neurodevelopmental disorders

The observation that multiple neurodevelopmental disorder-associated CaMKII variants, including F98S, E183V, and I205N, failed to form condensates with GluN2B suggests that LLPS is a conserved and essential property of CaMKII required for normal neuronal function. LLPS has been proposed to promote the local concentration of CaMKII and its substrates/binding partners at the postsynaptic density, thereby facilitating intermolecular phosphorylation and efficient signal propagation ^16,47,48^. Although these variants retained Ca^2+^/calmodulin-dependent kinase activity, they exhibited reduced condensate formation, consistent with the idea that LLPS enhances enzymatic efficiency within condensed assemblies. Thus, impaired LLPS likely compromises the self-sustaining activation of CaMKII and subsequent downstream signaling that underlies LTP and synaptic plasticity.

Importantly, neurodevelopmental disorder-associated CaMKII variants that failed to undergo LLPS also showed impaired interactions with a key synaptic partner, GluN2B, which normally anchor CaMKII at the PSD and link it to structural plasticity. This defect likely also disrupts association with additional proteins. Since CaMKII accumulation at excitatory synapses is critical for spine enlargement and stabilization, the disruption of LLPS could lead to impaired postsynaptic organization and weakened synaptic connectivity. More broadly, our results highlight LLPS as a fundamental molecular mechanism governing the assembly and regulation of postsynaptic signaling complexes ^15^. Similar to other condensate-forming synaptic proteins such as PSD-95, SynGAP, and SHANK3 ^49,50^, CaMKII LLPS enables the spatiotemporal coordination of synaptic signaling events in response to neuronal activity. Disruption of condensate formation may thus represent a unifying mechanism through which diverse molecular defects converge on synaptic disorganization and cognitive disruption in neurodevelopmental disorders.

Consistent with this view, our CaMKIIα I205K knock-in mice exhibited hyperactivity, reduced working and aversive memory, and abnormal social behaviors – phenotypes reminiscent of those observed in human neurodevelopmental disorders, including attention-deficit/hyperactivity disorder (ADHD). These findings suggest that LLPS-dependent organization of CaMKII is indispensable for both synaptic and behavioral homeostasis ^51,52^. Whereas the I205N individual was diagnosed with ADHD, the previously published F98S and E183V individuals both were still young at the time of publication, and no subsequent clinical follow-up has been reported to determine whether ADHD-like symptoms developed later ^33^. Nevertheless, ADHD is a common comorbidity across multiple neurodevelopmental disorders, raising the possibility that similar behavioral features may also occur in affected individuals.

### Perspective

How disruption of CaMKII LLPS and the resulting impairment in synaptic plasticity gives rise to hyperactivity remains unclear. Future studies should explore whether restoring LLPS can rescue synaptic or behavioral phenotypes in phase separation-deficient mutants. Structural or small molecule interventions that stabilize CaMKII-GluN2B interaction or enhance multivalent assembly at the PSD could potentially normalize aberrant signaling and plasticity. Moreover, the use of super-resolution imaging or *in vivo* biosensors could clarify how LLPS dynamics are modulated by neuronal activity and contribute to circuit-level plasticity. Together, our findings not only establish a direct mechanistic link between CaMKII LLPS behavior and synaptic organization, but also point to LLPS as a promising conceptual and therapeutic target for neurodevelopmental disorders.

## Acknowledgements

We thank laboratory members for technical assistance and discussion, and Professor Fukazawa (Fukui University) for providing the anti-GluA1–3 antibody. We thank Drs. Steven Middleton and Nozomi Asaoka for their comments on the manuscript. We are grateful for the help and support provided by the Integrative Bioscience Facility at Institute of Science Tokyo. The Airyscan confocal microscope used in this study was supported by MEXT Project for Promoting Public Utilization of Advanced Research Infrastructure (Program for Supporting Construction of Core Facilities; Grant Number JPMXS0440200025). This research was supported by JST CREST under JPMJCR20E4, AMED under JP24zf0127010, by JSPS under 18H04733, 18H05434, and 16H02455, HFSP research grant RGP0020/2019 to YH, 18K19377, 21H02595, and 21H05692 to TS, 24KJ1449 to RY, and by AMED 25wm0625123h0002 to YH and TS. Takeda Science Foundation Research Grants (2020), Research Grant from the Naito Foundation (2024), and Brain Science Foundation (2024) to TS. The Dutch Brain Organization (Hersenstichting, DR-2023-00429) and Stichting de Mere to GW, DV, and AJ.

---

**Figure S1.**
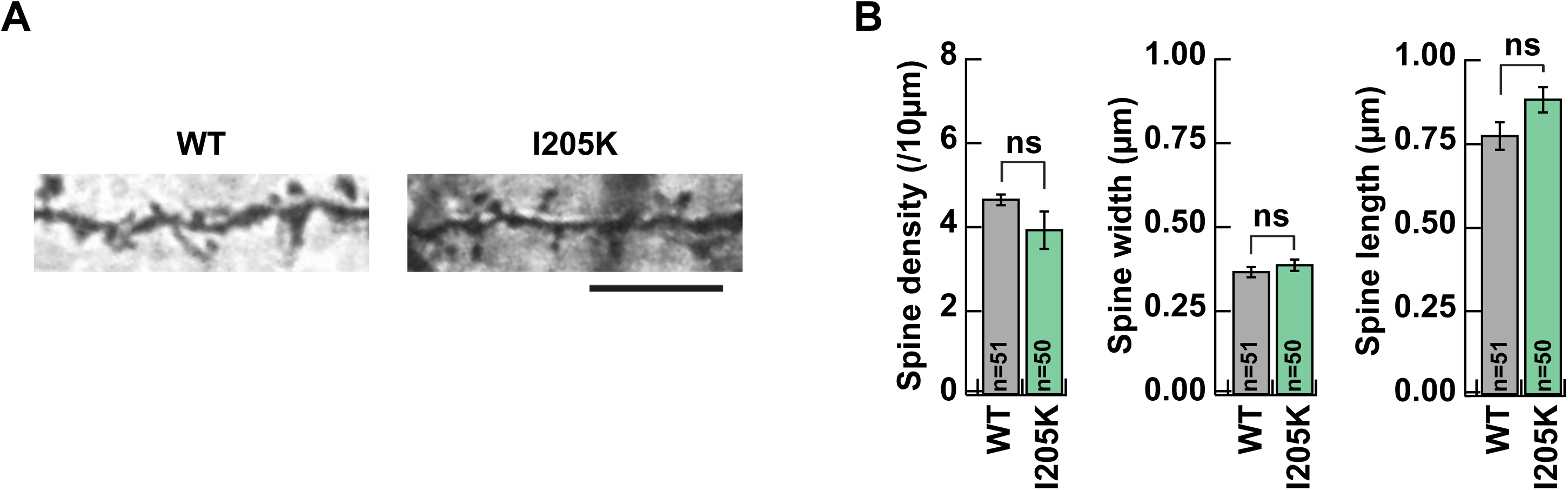
Spine morphology of CA1 pyramidal neurons in CaMKII I205K knock-in mice, related to Figure 2. A. Representative Golgi-Cox stained images of CA1 pyramidal neurons from WT and I205K mice. Scale bar, 10 μm. B. Quantification of spine density, head width, and spine neck length. (WT, n=51; I205K, n=50 neurons from 3 mice per genotype). Data are shown as mean ± SEM. ns, not significant, in a two-tailed unpaired t-test, compared to WT.

**Figure S2.**
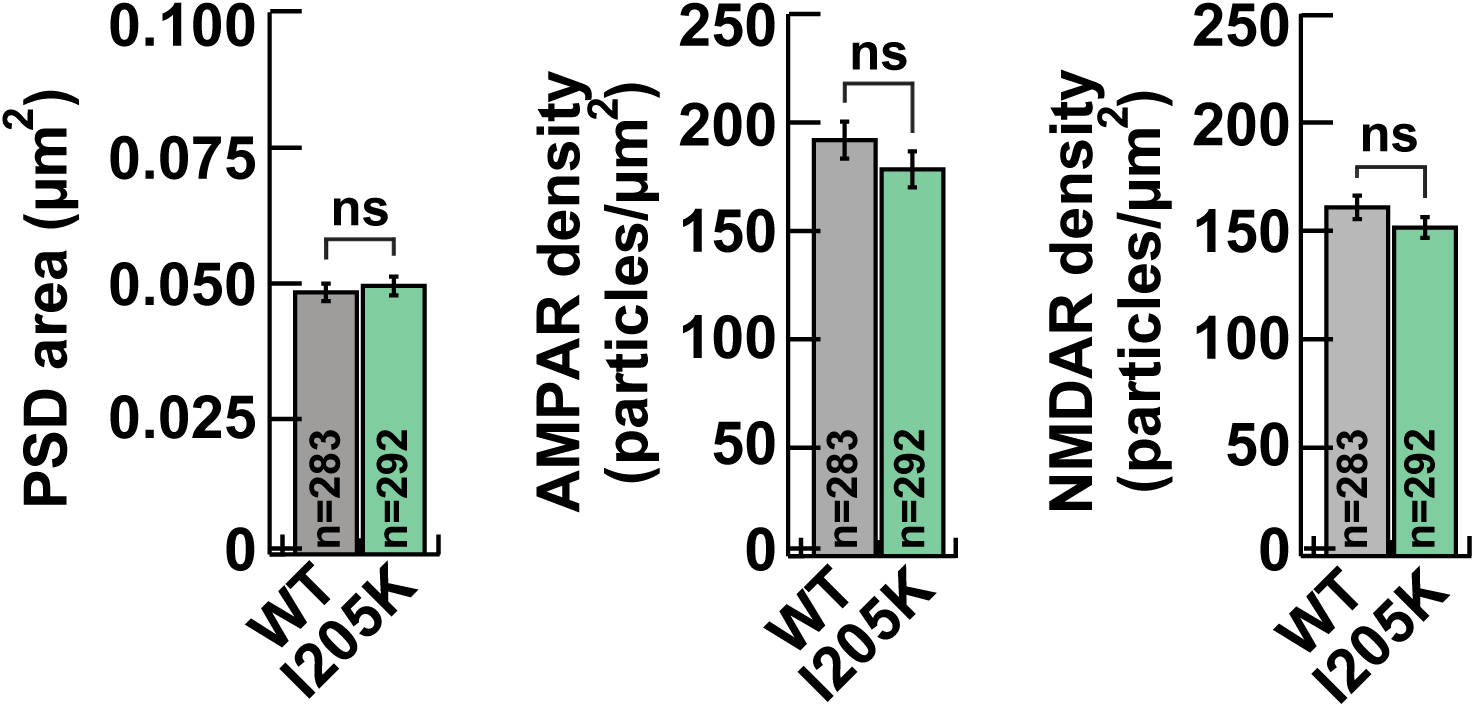
Comparison of PSD area and receptor density of spines in WT and CaMKII I205K knock-in mice, related to Figure 2. Quantification of PSD area, AMPAR and NMDAR density as labeled by immunogold particles (WT, n=283; I205K, n=292 synapses). Data are shown as mean ± SEM. ns, not significant, in a two-tailed unpaired t-test, compared to WT.

**Figure S3.**
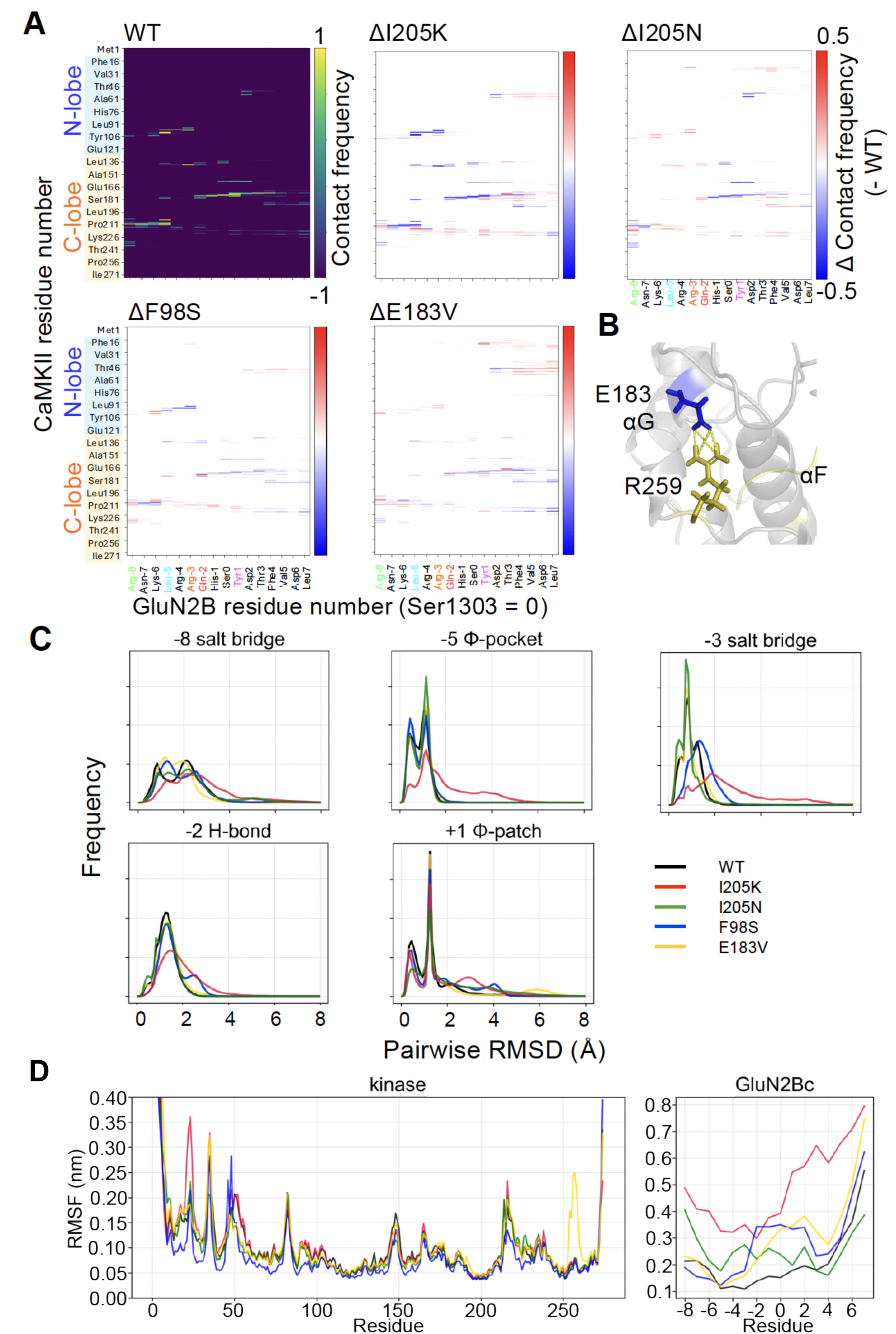
Contact map and structural dynamic analyses of WT and variant CaMKII-GluN2B complexes, related to Figure 7. A. Normalized contact maps of the GluN2Bc peptide and the kinase domain in WT (top left), along with the differences observed in each variant. In the difference maps, regions where interactions have increased are shown in red, and those where interactions have decreased are shown in blue. B. Interaction sites between E183 (blue) and R259 (yellow). C. Normalized RMSD distributions in WT (black), I205K (red), I205N (green), F98S (blue), and E183V (yellow) in each interaction site. D. Normalized RMSF in WT (black), I205K (red), I205N (green), F98S (blue), and E183V (yellow) in kinase domain (left) and GluN2Bc peptide (right). Both RMSF are calculated by aligning with backbone structures of the kinase domain.

**Table S1.** Patient information, related to Figure 6.

| FEATURE | VALUE |
| --- | --- |
| Gene altered | CAMK2A |
| Variant observed* | c.614T>A |
|  | p.Ile205Asn |
| Functional test |  |
| Pathogenic | yes |
| Direction of pathogenicity | Gain of Function |
| Mode of inheritance | de novo |
| Method of mutation detection | TRIO WES |
| Gender | male |
| Age at last investigation | 13 years |
| Siblings | 1 unaffected younger brother |
| Obstetric & Birth history |  |
| Spontaneous pregnancy | yes |
| Obstetric complications | mild hypertension with proteinuria and mild intrauterine growth restriction towards the end of gestation. |
| Labour spontaneous / induced | induced |
| Type of delivery vaginal, caesarian | vaginal |
| Gestational week | 39 |
| Birth weight (grams/SD) | 2730 (-1.75 SDS) |
| Birth length (cm/SD) | 49cm (-0.56 SDS) |
| Birth head circumference (cm/SD) | 33cm (-2 SDS) |
| Asfyxia | no |
| Apgar score | 9/10 |
| Birth complications yes/no | no |
| Discharge hospital within 48 hours yes/no | yes |
| Growth |  |
| Growth abnormality yes/no | no |
| Height at last investigation (cm/SD) | 162cm (+0.19 SDS compared to target height) |
| Weight at last investigation (kg/SD) | 40.7kg (-1.48 SDS weight for height) |
| Head circumference at last investigation (cm/SD) | 53.5cm (-0.76 SDS) |
| Developmental History |  |
| Degree of intellectual disability | mild |
| Age of rolling | 5 months |
| Age of independent sitting | 9 months |
| Age of independent walking | 19 months |
| Equipment for mobilisation | no |
| Abnormal tone | generalized hypotonia |
| Muscle weakness | no |
| Involuntary movements | no |
| Current Level Gross Motor Functioning | able to run and walk stairs without problems, plays soccer and judo. Movement ABC 2: total test score 20, percentile score 0.1 (delayed motoric development) |
| Coordination/balance difficulties | no (resolved) |
| Current level Fine motor functioning: | writes in big letters, not able to tie shoelaces. Manual dexterity M-ABC-2: standard score 1, percentile 0.1 |
| Potty-trained for stool/urine | yes (age 2.5 years) |
| Independent eating | yes |
| Independent clothing | yes |
| Age of first words | unknown (delayed) |
| Level of understanding of spoken language | yes, two step commands |
| Level of speech | verbal, can make long sentences. Dysarthria and syntax problems. |
| Equipment for communication | no |
| **Regression or stop in development** | no |
| **Neurology** |  |
| **Seizures** | no |
| **EEG anomalies** | yes, frequent focal epileptiform abnormalities (sharp waves and spike-and-slow wave complexes) right fronto-temporal |
| **Autonomic dysfunction** | no |
| **Sleeping problems** | no |
| **Brain imaging performed** | not performed |
| **Structural abnormalities Organs** |  |
| **Respiratory anomalies** | no |
| **Cardiac anomalies** | no (physiological systolic murmur) |
| **Upper Gastro-intestinal abnormalities** | no |
| **Swallowing problems** | no |
| **Drooling** | no |
| **Lower Gastro-intestinal abnormalities** | no |
| **Equipment for feeding** | no |
| **Urogenital/kidney anomalies** | unknown |
| **Skin abnormalities** | no |
| **Joint hypermobility** | yes |
| **Visual impairment** | no |
| **Hearing impairment** | no |
| **Skeletal abnormalities** | no |
| **Anemia** | no |
| **Immune system abnormalities** |  |
| **Recurrent, atypical or severe infections** | no |
| **Laboratory abnormalities** | no |
| **Behavioural anomalies** |  |
| **Atypical behaviour core to phenotype** | yes |
| **DSM-V classification** | ADHD diagnosis |
| **Pervasive Symptoms** | no |
| **ADD/ADHD Symptoms** | yes |
| **Mood disorder symptoms (depressive, anxiety)** | situational anxiety appropriate for developmental age |
| **Bi-polar disorder symptoms** | no |
| **Self-harm** | no |
| **Clinically recognisable dysmorphisms** |  |
| **Facial** | no clear dysmorphisms, eyes mildly deep-set, subtle upslant |
| **Thorax abnormality** | mild high thoracic left-convex scoliosis |
| **Hands** | no |
| **Feet** | clinodactyl dig V |
| **Other** |  |
| **Other anomalies or problems** | no |
| **Additional genetic findings** | no |

## Notes

### Competing Interest Statement

The authors have declared no competing interest.

